# Causes and biophysical consequences of cellulose production by *Pseudomonas fluorescens* SBW25 at the air-liquid interface

**DOI:** 10.1101/538561

**Authors:** Maxime Ardré, Djinthana Dufour, Paul B Rainey

## Abstract

Cellulose over-producing wrinkly spreader mutants of *Pseudomonas fluorescens* SBW25 have been the focus of much investigation, but conditions promoting the production of cellulose in ancestral SBW25, its effects and consequences have escaped in-depth investigation through lack of in vitro phenotype. Here, using a custom built device, we reveal that in static broth microcosms ancestral SBW25 encounters environmental signals at the air-liquid interface that activate, via three diguanylate cyclase-encoding pathways (Wsp, Aws and Mws), production of cellulose. Secretion of the polymer at the meniscus leads to modification of the environment and growth of numerous micro-colonies that extend from the surface. Accumulation of cellulose and associated microbial growth leads to Rayleigh-Taylor instability resulting in bioconvection and rapid transport of water-soluble products over tens of millimetres. Drawing upon data we build a mathematical model that recapitulates experimental results and captures the interactions between biological, chemical and physical processes.

**IMPORTANCE:** This work reveals a hitherto unrecognized behaviour that manifests at the air-liquid interface, which depends on production of cellulose, and hints to undiscovered dimensions to bacterial life at surfaces. Additionally, the study links activation of known diguanylate cyclase-encoding pathways to cellulose expression and to signals encountered at the meniscus. Further significance stems from recognition of the consequences of fluid instabilities arising from surface production of cellulose for transport of water-soluble products over large distances.

## INTRODUCTION

Surfaces are frequently colonised by microbes. Surface-associated microbes grow as dense populations / communities termed “biofilms” (1, 2, 3). Growth at surfaces provides microbes with nutrients and opportunities for cross-feeding (4, 5). For pathogens, surface colonisation is often a prelude to invasion (6, 7). Microbes in high-density populations can find protection against external factors such as antibiotics and toxic agents (8). At the same time, microbes in biofilms experience intense competition for resources and can be negatively impacted by costs associated with exposure to metabolic waste products (9). For long-term survival, escape from surfaces and dispersal is crucial (10).

Primary attention has been given to colonisation of solid-liquid surfaces (11, 12). This owes as much to the importance of these surfaces as it does the ease with which they can be studied. For example, colonisation of abiotic surfaces can be measured by simple histochemical assay or by microscopic observation using flow cells (13, 14). Decades of study have revealed insight into the role of adhesive factors including polymers and proteinaceous adhesions involved in surface attachment and the regulatory pathways controlling their expression (15). A particular focus has been pathways for synthesis and degradation of the secondary signalling molecule cyclic-di-GMP (16). For the most part, the precise signals activating these regulatory pathways are unclear. Moreover, the frequent use of mutants — sometimes intentionally, but often inadvertently — that constitutively over-produce adhesive factors has stymied progress in understanding many subtleties surrounding surface colonisation.

Surfaces are also a feature of the interface between gas and liquid, but colonisation of such surfaces has been received much less attention (17, 18, 19, 20). Air-liquid interfaces (ALIs) are of special relevance for aerobic organisms because colonisation of the meniscus provides access to oxygen. While many motile aerobic bacteria display taxis toward oxygen, this alone is often insufficient to allow cells to overcome the effects of surface tension necessary to colonise the ALI. Where colonisation is achieved, in the absence of mechanisms promoting buoyancy, cells must contend with the effects of gravity that become increasingly challenging with build up of biomass.

The interface between air and liquid has further significance in that it often marks the divide between aerobic and anaerobic conditions. This has implications for surface chemistry with ensuing physiological effects for bacteria. For example, iron, an essential element, exists in the insoluble and biologically unavailable ferric form in the presence of oxygen, but is water soluble and freely available in the absence of oxygen (21). Bacteria growing within an initially resource-rich and oxygen replete broth phase consume oxygen and thus further growth requires access to the ALI (22). Bacteria that achieve colonisation of this surface must then contend with iron deplete conditions requiring the synthesis of siderophores (23).

To date, studies of colonisation of the ALI have been largely centred on genotypes that constitutively produce polymers such as cellulose (24). Often these have arisen as a consequence of selection experiments in static broth microcosms where mutants with constitutively active diguanylate cyclases (and ensuing constitutive production of the respective polymers) have a selective advantage that arises from capacity to form dense microbial mats (pellicles) at the ALI (25, 26, 27, 20). While such mutants have made clear the central importance of cellulose and related polymers (28), the generality of conclusions arising from the use of constitutively active mutants need to be treated with caution (26). Desirable would be analysis of the biophysics of ALI colonisation in wild type bacteria where regulation of polymer production is unaffected by mutation.

Almost two decades ago it was reported that in well-mixed culture the fitness of a cellulose-defective mutant of *Pseudomonas fluorescens* SBW25 was equivalent to that of the wild type (ancestral) bacterium (24). Also reported in that study was a significant reduction in fitness of a cellulose defective mutant in static broth culture, but the reasons were not determined. Recent observations of the growth of a cellulose-defective mutant of wild type (ancestral) SBW25 made during the course of analyses of evolutionary convergence in polymer production by SBW25 (28) led to the realisation of a subtle phenotype associated with absence of growth in the cellulose-defective mutant at the air-liquid interface. Unlike ancestral SBW25, the mutant grows exclusively within the broth phase with ensuing negative effects of oxygen limitation responsible for its previously noted low fitness (24).

Here we seek to understand the biological role of cellulose and do so via a device that combines spectrophotometry with multi-perspective time-lapse imaging. Aided by the device we monitor surface growth, reveal the contribution made by cellulose and show that it involves regulatory contributions from three known diguanylate cyclase-encoding regulatory pathways. The production of cellulose allows formation of a lawn of micro-colonies at the meniscus that eventually coalesce into a thin film of bacteria. The mass of bacteria and cellulose generates a gravitational force that leads to Rayleigh-Taylor instability and causes bioconvection (29). One consequence of bioconvection is the rapid transport of the water-soluble iron-binding siderophore, pyoverdin. A mathematical model based on partial differential equations with fluid dynamics described by the Navier-Stokes (NS) equation with Boussinesq approximation accounts for the diffusion-reaction and convection processes occurring in the microcosm.

## RESULTS

Describing microbial colonisation of the air-liquid interface (hereafter ALI), although in principle straightforward, is fraught with difficulty. While advanced microscopic techniques offer possibilities to observe colonisation at the single cell level, much stands to be gained from more macroscopic perspectives, aided by low power microscopy in conjunction with time-lapse photography.

### Device

To understand and measure growth of ancestral SBW25 and the cellulose-defective mutant SBW25 Δ*wssA-j* a device was constructed that allows growth at the ALI and in the broth phase to be monitored from multiple perspectives (Fig. 1). It comprises three cameras: one placed perpendicular to the microcosm to record growth within the microcosm and on the under surface of the meniscus, one mounted at a 45° angle above the ALI to capture surface growth and one to detect the light emitted from excitation of the fluorescent signal arising from production of the iron-chelating siderophore, pyoverdin. Additionally, the device incorporates a laser and corresponding photodiode to vertically scan the flask at regular (5 min) time intervals.

**FIG 1.**
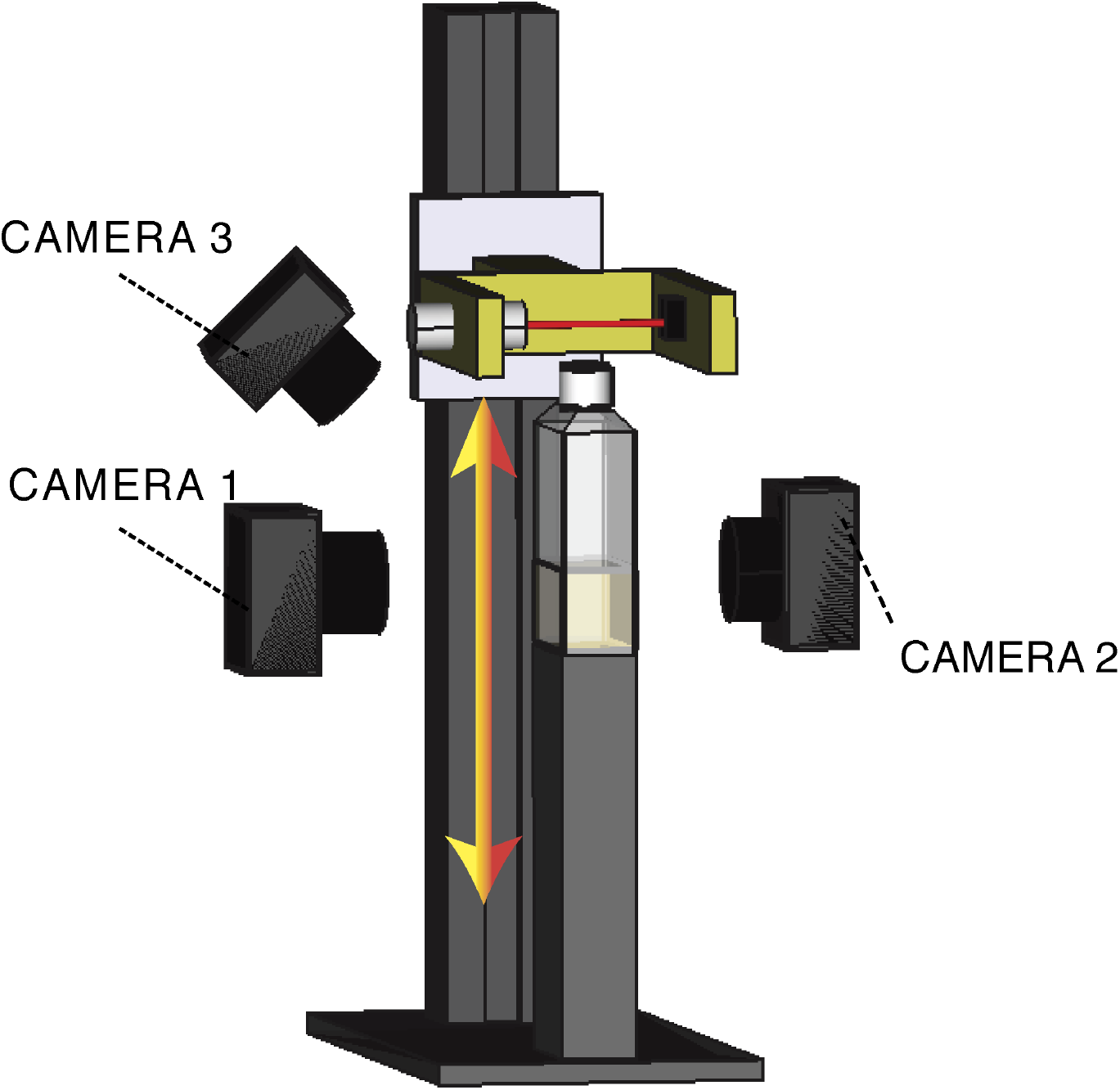
Experimental device. A polycarbonate cell-culture bottle filled with 20 ml of KB and inoculated with bacteria is placed on a fixed vertical stand. The device and associated cameras are maintained within a 28°C incubator. The flask is scanned vertically every 5min with a 600 nm laser beam with 1mm section. Light passing through the flask is collected by a photodiode. To obtain a measure of the optical density in the flask along a vertical profile, the alignment laser-photodiode is coupled to a motorised device that ensures smooth vertical translation. Three cameras are located around the f ask. The first (camera 1) obtains a side-view image of the liquid phase of the medium using bright-field illumination. The second (camera 2), also fixed perpendicular to the flask, monitors fluorescence associated with pyoverdin (excitation 405/emission 450 nm). The third camera (camera 3) is oriented with a 45° angle and captures growth at the ALI using bright-field illumination.

### Cellulose is required for colonisation of the ALI

Figure 2 shows the growth dynamics of ancestral SBW25 and SBW25 Δ*wssA-J* determined by the scanning laser and calibrated using direct plate counts. SBW25 Δ*wssA-J* is slower to enter exponential growth than SBW25, it grows at approximately the same rate (SBW25, 0.53+/−0.02 h-1; SBW25 Δ*wssA-J* 0.57+/−0.03 h^−1^), but density in stationary phase is consistently lower. Notable in SBW25 at 24 hours is a reproducible plateau of growth followed by a further increase and a widening of difference in cell density compared to SBW25 Δ*wssA-J* (Fig. 2). No such intermediate plateau occurs in the cellulose mutant.

**FIG 2.**
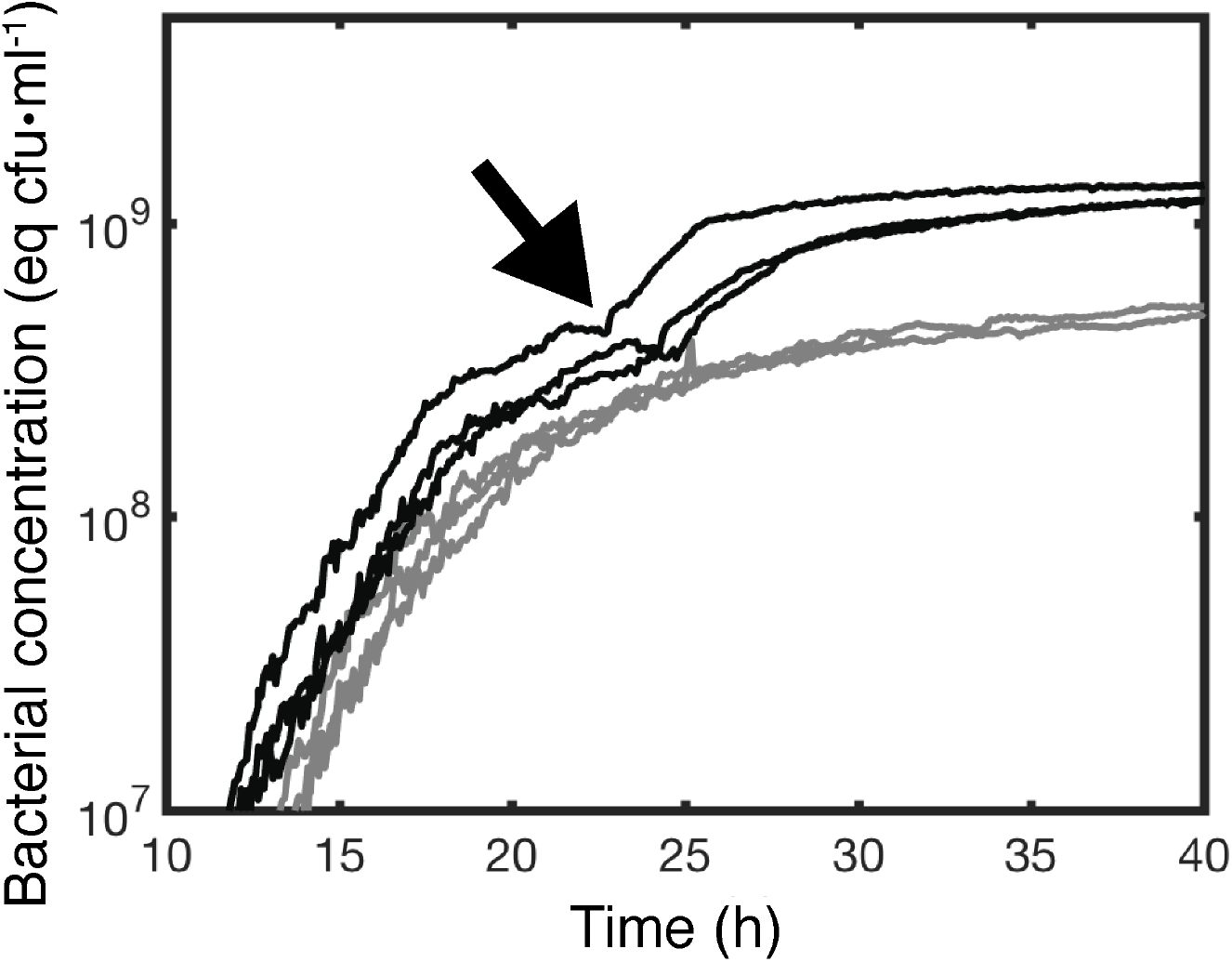
Production of cellulose maximises growth in static broth culture. Dynamics of growth of *P. fluorescens* SBW25 (black lines) and *P. fluorescens* SBW25 Δ*wssA-J* (cellulose negative mutant) (grey lines) in unshaken KB as determined by the scanning laser device and associated photodiode depicted in Figure 1. Every curves is an independent experiment made in a new flask. Data are spatial average of the optical density at 600 nm (0D600) obtained from scanning the vertical section of a flask. 0D600 measures are calibrated using direct plate counts of colony forming units (equivalent cfu·ml^−1^). Measurement were taken every 5min. The arrow denotes the onset of bioconvection caused by production of cellulose that marks a secondary increase in growth. This second growth phase is absent in the cellulose negative mutant.

Time-lapse observation of the ALI from a 45o angle in flasks inoculated with SBW25 reveal presence of a thin film at 19 h that is more prominent at 26 h and still evident albeit weakly at 40 h (Fig. 3a and supplementary movie file 1). Beyond the 40 h time period wrinkly spreader mutants arising within the flasks begin to grow at the ALI. In contrast, no evidence of colonisation of the ALI is evident in SBW25 Δ*wssA-J* (Fig. 3b and supplementary movie file 2). Observations from the camera perpendicular to the flask confirmed presence of surface growth in SBW25 (Fig. 3a), but not in the cellulose mutant (Fig. 3b). Additionally rapid streaming was observed in the broth phase for the ancestral genotype but not for SBW25 Δ*wssA-J* (Fig. 3a and 3b, supplementary movie files 3 and 4 respectively). The significance of this streaming dynamic is considered in detail below.

**FIG 3.**
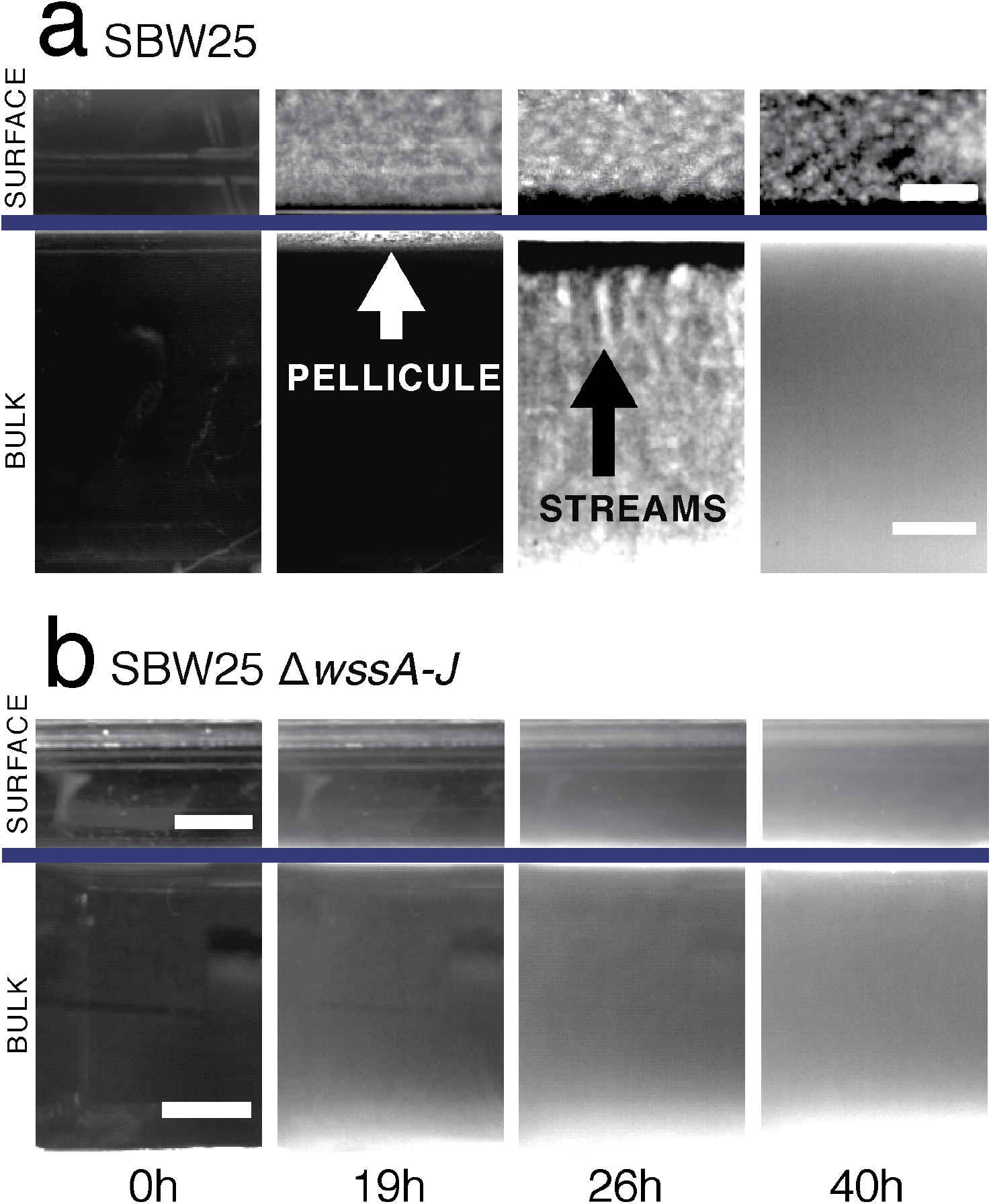
Cellulose is necessary for growth at the air-liquid interface (ALI) and results in bioconvection. Bright-field images of ancestral *P. fluorescens* SBW25 (a) and *P. fluorescens* SBW25 Δ*wssA-J* (cellulose negative mutant) (b) taken at four time intervals. Complete movies are available as SI movie files 1,2,6 and 7. Images above the solid line show growth at the ALI captured using camera 3; images below the line are from camera 1 (see Figure 1). At time 0 h the medium is inoculated with −10^4^ cells·ml^−1^. By 19h the ancestral cellulose-producing genotype has formed a thin white pellicle at the ALI (visible by both camera 1 and 3). No pellicle formation is seen in the cellulose negative mutant, but growth is evident in the broth phase. By 26 h, in cultures of the cellulose-producing ancestral type, plumes characteristic of bioconvection stream from the ALI (pointed by the black arrow). No evidence of mat formation or streaming is seen in SBW25 Δ*wssA-J.* By 40h streaming has largely ceased in the ancestral type, although growth is still apparent at the ALI. Scale bars are 5mm. Contrast has been adjusted to highlight salient features.

Curious as to the nature of the previously unseen surface growth we obtained high-resolution photos at hourly intervals (between 15h and 20 h) from directly above the surface using a light source for illumination positioned at an oblique angle to the surface. No surface growth was evident for SBW25 Δ*wssA-J* (Fig. 4b), but remarkably, from the ancestral genotype, numerous micro-colonies emerged from the surface of the meniscus and grew outward as if on an agar plate (Fig. 4a). By 19 hours microcolonies can be seen to fall from the surface through the effects of gravity, but is quickly followed by coalescence and collapse of the entire population of microcolonies (Fig. 4a and supplementary movie file 5). Interestingly, at the moment of coalescence and mat collapse “chewing gum-like” strands suddenly appear at the ALI, which is more characteristic of standard pellicles (20, 24, 30). This raises the possibility that cellulose is transformed from a viscous liquid to a solid by the stretching effect of gravity.

**FIG 4.**
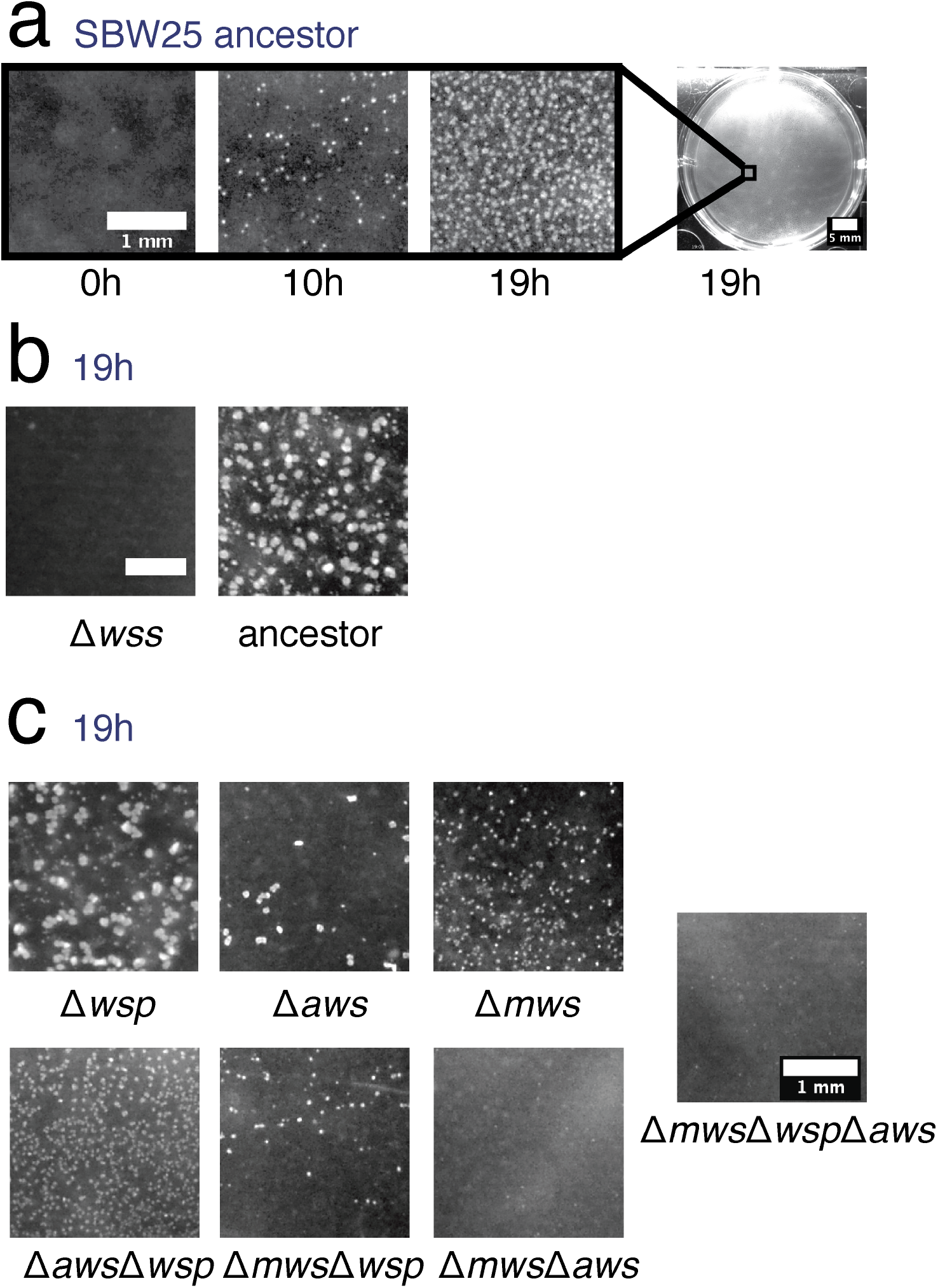
Multiple diguanylate cyclase are required for colonisation of the ALI. Microcolony formation at the ALI for ancestral *P. fluorescens* SBW25 and a range of mutants captured from a camera mounted directly above individual wells of a six-well tissue culture plate containing 5ml KB. Time course of micro-colony formation for ancestral *P. fluorescens* SBW25 (a). Comparison with SBW25 Δ*wssA-J* (cellulose negative mutant) at 19h (b). Patterns of micro-colony formation at 19 h in mutants devoid of Wsp (Δ wsp), Aws (Δαws) and Mws (Δmws) diguanylate cyclase-encoding pathways and combinations there-of. Scale bar is 1 mm, except for the entire well in (a) which is 5mm.

### Regulation of cellulose and ALI colonisation by multiple diguanylate cyclase-encoding regulatory pathways

Numerous studies of constitutive cellulose overproducing mutants — the so named wrinkly spreader (WS) types (31) — have shown the phenotype to arise primarily by mutations in the Wsp, Aws and Mws pathways (27, 28, 29, 31, 32, 30). Mutations in the negative regulators of these diguanylate cyclase-encoding pathways (DGCs) result in over-production of cyclic-di-GMP, overproduction of cellulose and formation of substantive and enduring mats at the ALI. While these findings have connected over-expression of DGC-encoding pathways to the WS phenotype, the relationship between known DGC-encoding pathways and cellulose expression in the absence of DGC over-activating mutations has been a mystery. Recognition that ancestral SBW25 activates cellulose production at the ALI leading to micro-colony formation and a frail film of cells, allowed investigation of the role of Wsp, Aws and Mws in expression of this phenotype.

A reduction in the formation of micro-colonies in SBW25 Δ*wspABCDEFR*, SBW25 Δ*awsXRO*, and SBW25 Δ*mwsR* demonstrates for the first time a connection between the Wsp, Aws and Mws pathways, the production of cellulose and colonisation / microcolony formation at the ALI (Fig. 4c) in ancestral SBW25. Surprisingly, no single pathway mutant resulted in a cellulose defective phenotype that matched that of the cellulose defective *wssA-J* deletion mutant (Fig. 4c). Equally surprising was that all three pathways make some contribution to colonisation of the ALI (Fig. 4c). The most pronounced phenotype was associated with SBW25 Δ*mwsR,* followed by SBW25 Δ*awsXRO* and SBW25 Δ*wspABCDEFR.* A mutant lacking all three pathways was indistinguishable from SBW25 Δ*wssA-J* (Fig. 4c).

### Cellulose causes bioconvection

As noted above, in microcosms inoculated with cellulose-producing ancestral SBW25, material falls in finger-like plumes that stream from the ALI (Fig. 3a). Analysis of time-lapse movies (supplementary movie file 6) shows plumes to be characteristic of long-range convection (Fig. 5), which arises as a consequence of instability of the interface between the cellulose-rich meniscus and the less dense broth phase beneath. The phenomenon is known as Rayleigh-Taylor instability. That cellulose is the critical component stems from the fact that the streaming plumes are evident in ancestral SBW25, but not in cultures of the cellulose negative mutant (SBW25 Δ*wssA-J*).

**FIG 5.**
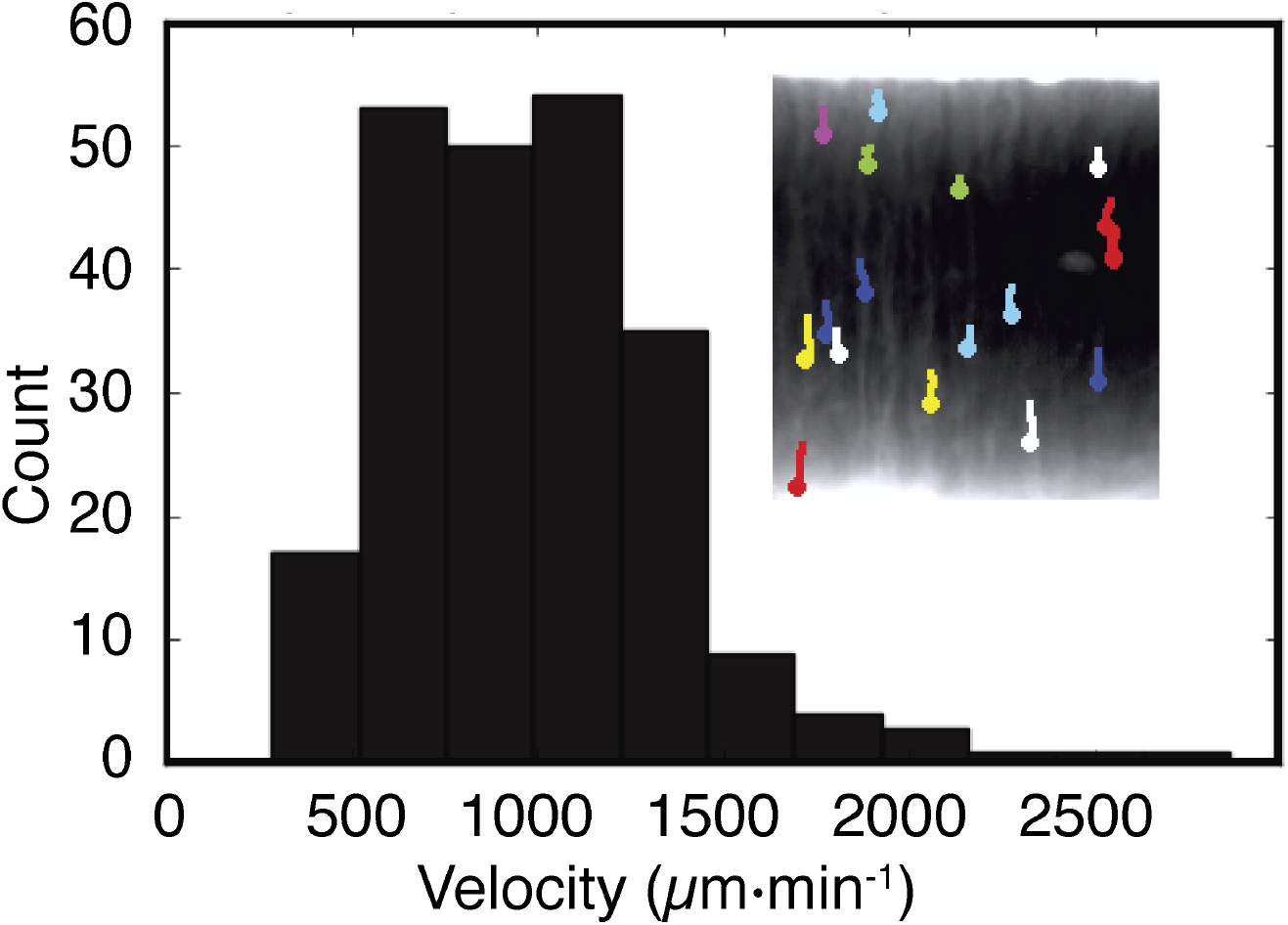
Bioconvection caused by cellulose. Time lapse images via bright field camera 1 (Fig. 1) capture biomass dynamics in the liquid medium. By 25 h Rayleigh-Taylor instability generates plumes of biomass that fall from the ALI to the bottom of the flask (inset). The velocity of movement is obtained by tracking trajectories of the plumes. The frequency distribution of plume velocity reveals a mean speed of 983 ± (SD) 373 μm·min^−1^.

Quantification of the streaming plumes shows instability at ~25h and continues until ~ 40h at which point streaming ceases and the medium becomes homogeneous. The velocity of the falling plumes ranges from 500 to 2000μm·min^−1^ (Fig. 5). From this it is possible to calculate the Péclet number that defines the contribution of diffusion relative to bioconvection on the transport of water-soluble products. In this instance the Péclet number (Pe) is ~1000 (calculated by multiplying the typical plume length (1cm) by its velocity (~1 ·10^−3^cm·s^−1^) and then dividing by the diffusion coefficient of pyoverdin −1 ·10^−6^cm^2^· s^−1^). The Péclet number, being greater than 1 (Pe is a dimensionless number), means that bioconvection is a more significant contributor to the transport of soluble products than diffusion.

### Bioconvection affects spatial distribution of extracellular products

A soluble product of relevance to *P. fluorescens* SBW25 in static culture is the water-soluble iron binding siderophore, pyoverdin (33). That it is fluorescent means that it is readily monitored. Figure 6a shows the average concentration of pyoverdin at the ALI as imaged via camera 2 equipped with suitable optical filters (see Fig. 1). The first indication of pyoverdin production occurs at the ALI at ~ 19h and coincides precisely with the first visible stages of surface colonisation where micro-colonies begin to form at the meniscus (Fig. 4).

**FIG 6.**
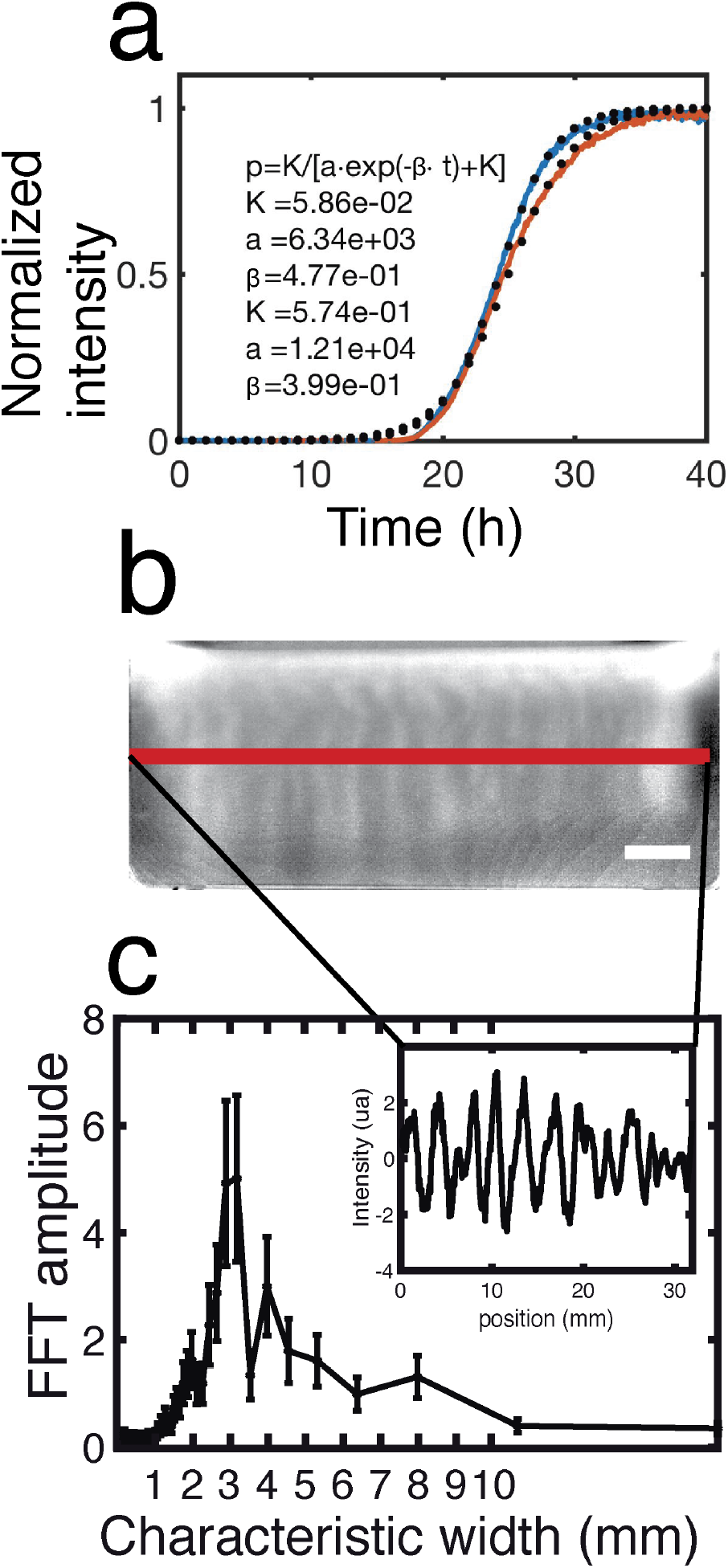
Camera 2 (see Figure 1) monitors pyoverdin concentration in the flask by measuring fluorescence. Pyoverdin is produced primarily at the ALI. The average fluorescence along the ALI increases with time like a sigmoid curve (*a*). The adhoc logistic function of the inset gives the normalized intensity (*p*) as a function of the time (*t*) and the parameters of the fit (*K, a* and *β*). The fitted curves (dotted line) adjust the experimental curves (plain line) for the estimated values of the parameters given in inset. Plumes due to Rayleigh-Taylor instability transport pyoverdin from the ALI to the liquid phase. Pyoverdin concentration is transiently higher along vertical columns that correspond to the plumes flowing from the ALI. The white scale bar is 5mm. The fluorescence intensity profile along the red horizontal line (b) shows that pyoverdin is distributed with a fluctuating spatial structure (inset). Fast Fourier transformation (FFT) of the intensity profile reveals these fluctuations to have a characteristic wavelength of 3mm.

The first signs of pyoverdin production are restricted to the ALI despite the fact that at 19h and thereafter, the broth phase is turbid with growth (supplementary movie file 3 shows turbidity in the flask and supplementary movie file 7 shows pyoverdin in the flask). This is consistent with oxygen being available at the broth surface (and absent in the bulk phase due to metabolic activity) causing iron at the ALI to exist in the insoluble ferric form, leading to activation of pyoverdin synthesis solely at the ALI. The kinetics of pyoverdin production were quantified by fitting data to a simple logistic model (Fig. 6a) whose fit indicates that the underlying chemical reaction is autocatalytic and characteristic of positive feedback regulation that controls pyoverdin synthesis (33).

Visible plumes of pyoverdin (Fig. 6b) were quantified by measuring pixel intensity across a single horizontal profile (inset Fig. 6c) as indicated by the red line in Figure 6b. To determine the characteristic plume width (Fig. 6c), the data were analysed by Fast Fourier transformation (FFT). The transformation shows that pyoverdin is concentrated in plumes with a horizontal width of 3mm.

### Modelling microcosm dynamics

Surface colonisation by *P. fluorescens* SBW25, interaction of cells with oxygen and ensuing effects, including bioconvection and transport of pyoverdin, draw attention to striking ecological complexity in this simplest of microcosms. To determine the match between current understanding of the interplay between biological, chemical and physical processes and the extent to which simple biophysical mechanisms explain the observed dynamics, we constructed a model based on diffusion-reaction processes and hydrodynamics. The degree of fit between model and data stands to show how well the system is understood.

The model is based on experimental quantification of bacterial culture density, pyoverdin concentration, and fluid flow. It uses partial differential equations to account for the diffusion-reaction-convection processes within the flask. The local concentration of bacteria, oxygen, pyoverdin and cellulose are described as continuous fields. The liquid environment is modelled as an incompressible Newtonian fluid with a mass density that depends on the concentrations of bacteria and cellulose. Its dynamic is described by the Navier-Stokes (NS) equation using the Boussinesq approximation, in which the variations of density are neglected except in the buoyancy force (34). The coupled equations allow for inclusion of different physical interactions between the components. Details are provided in the Materials and Methods section.

The model was solved numerically as a means of validation. Simulations were performed on a two-dimensional grid representing a physical domain of size 1 cm2. The top of the domain corresponds to the ALI with free fluid slip (liquid can move along the ALI) and no penetration boundary conditions (the meniscus cannot be deformed). The sides correspond to the lateral walls of the microcosm and the bottom of the flask. The boundary conditions on the wall allow no fluid slip (liquid cannot move along the wall) and no penetration.

The results of the simulation are shown in Figure 7 (and supplementary movie files 8-11) and closely reproduce the dynamics observed in microcosms. Bacteria replicate and consume oxygen until growth saturates at ~ 3 · 10^8^cfu·ml^−1^. At I6h oxygen is available at the meniscus and in a single millimetre layer immediately below the ALI. Also at I6h pyoverdin production begins; at I9h the first indication of cellulose production become visible resulting in an increase in density of the surface layer. Soon after, cellulose-laden regions begin to form descending plumes marking the onset of Rayleigh-Taylor instability. Plumes flow from the ALI to the bottom of the flask at a speed of~1000μm·min^−1^. This is in accord with experimental observations. Additionally, plumes serve to transport pyoverdin (over a millimetre scales) and oxygen, which penetrates several millimetres into the liquid phase. Robustness of the model to changes in parameter settings was assessed by performing six simulations over a range of parameter values. Changes to *c*_0_ and *o*\* made minimal difference over multiple orders of magnitude. Changes of one order of magnitude in the values of *b*\* and *ρ_c_* eliminated bioconvection, which is expected given that these parameters are directly proportional to the mass term in the Navier-Stokes equation. Alterations to parameters *V*\* and *γ* changed the dynamics of the system leading to a delay in the onset of bioconvection. The results are shown in supplementary data file “supplementaryFile12”.

**FIG 7.**
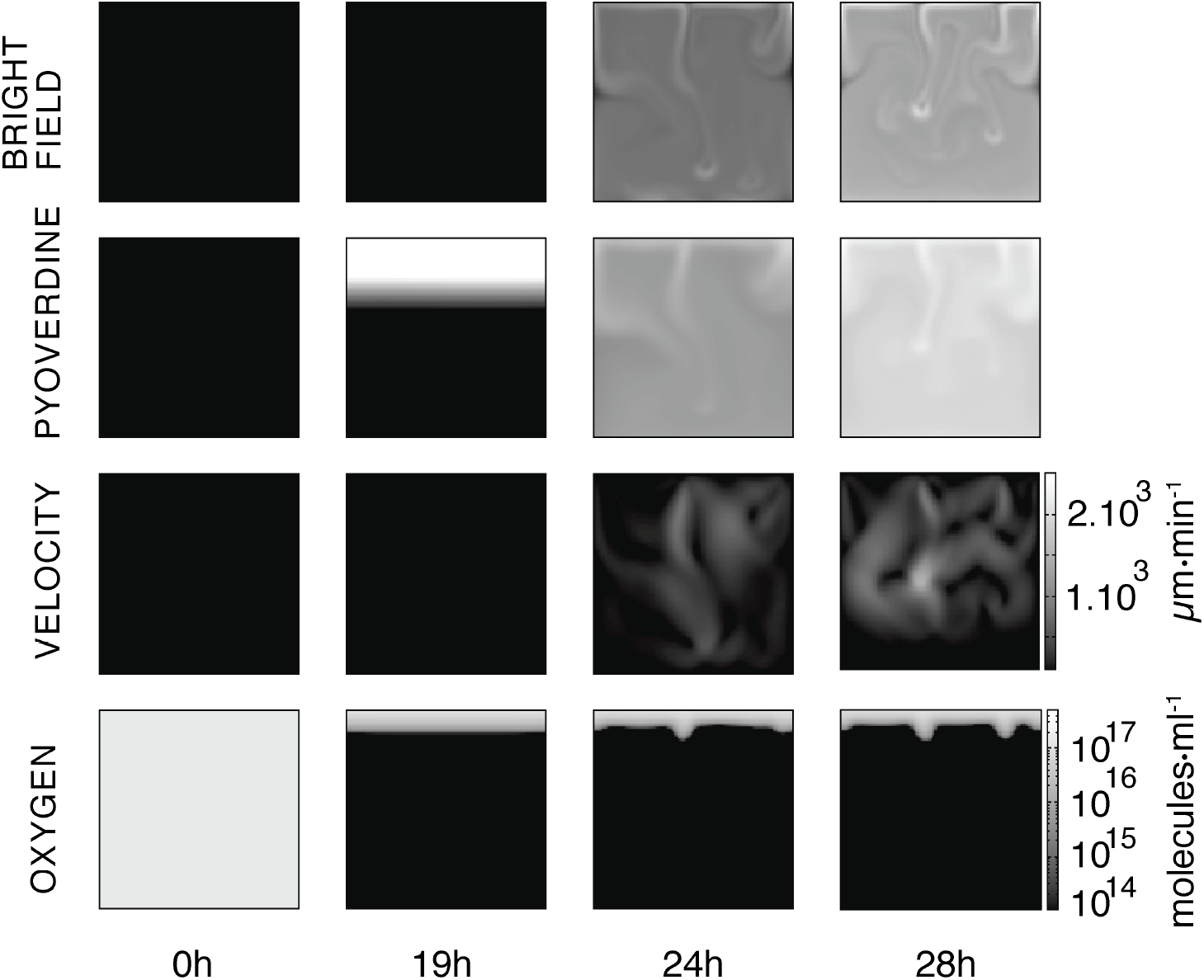
Numerical simulation of the mathematical model. Images display the dynamics of the simulated microcosm from inoculation at 0h to 28h. Time resolved movies are available in supplementary movies 8-11. The first row above shows the dynamics of the biomass in the bulk (bacteria and cellulose) as if observed with bright field illumination 1 (Fig. 1). In experiments, at 24h, plumes concentrated in biomass flow are evident in the liquid phase. The second row shows the concentration of pyoverdin in the liquid phase. The plumes transport pyoverdin into the bulk phase. The third row shows the dynamics of liquid velocity. When bioconvection is activated fluid flow is of the order of 1000 μm·min^−1^, which is consistent with the measurements shown (Fig. 5). The fourth row shows the dynamics of oxygen concentration. Soon after inoculation oxygen in the bulk phase is eliminated due to metabolic (oxygen consuming) activities of bacteria. The supply of oxygen at the ALI combined with growth of bacteria and production of cellulose means a gradient of oxygen 2-3 mm into the liquid. Images at 24h and 28h show that oxygen transport from the ALI before consumption by bacteria in the liquid phase. The square images are 1 cm2 and contrast is identical across each row.

## DISCUSSION

The interface between liquid and air defines a niche of significance for many bacteria (4). For aerobic organisms it is an environment replete with oxygen, it offers opportunities for unfettered surface spreading that may aid dispersal and indirectly, may allow rapid colonisation of solid surfaces; colonisation of the ALI may also allow bacteria to escape grazing by solid-surface associated predators. Despite its ecological relevance knowledge of mechanisms and consequences of surface colonisation are poorly understood.

For more than two decades studies of evolution in experimental microcosms have drawn attention to adaptive mutants of *P. fluorescens* SBW25 that specialise in colonisation of the ALI (24,25,31,35). These mutants, which constitutively overproduce cellulose as a consequence of DGC-activating mutations (26, 27, 28, 32, 30, 36), reap a significant adaptive advantage in static broth microcosms because of ability to grow at the ALI and thus access to oxygen. Largely unknown however has been the ecological significance of cellulose in the ancestral type and more generally, the role of cellulose in the natural environment. Impeding progress has been the fact that cellulose production is not evident on standard agar plate culture and neither is it produced in shaken broth culture. In the absence of a phenotype in vitro it is difficult to make progress.

Nonetheless, several previous studies have indicated environmental relevance: Gal et al (37) showed a cellulose defective mutant to be significantly less fit than the ancestral type in assays of plant colonisation and Giddens et al (38) showed the cellulose-encoding wss operon to be specifically activated on plant root surfaces. Koza et al showed that addition of metals including iron and copper to KB caused induction of a mucoid cellulose-containing agglomeration at the ALI (39). Perhaps the most significant finding, but at the time overlooked, was from competitive fitness assays between ancestral SBW25 and a wss defective mutant performed in shaken and unshaken microcosms (24): in shaken culture the fitness of the cellulose defective mutant was no different to the ancestral type, but in unshaken culture the mutant was significantly less fit. Here, prompted by recent observation of the poor growth in unshaken culture of SBW25 Δ*wssA-J* (28), combined with new tools of observation, we have come a step closer to understanding the biological significance of bacterial cellulose production.

Apparent from use of the device shown in Figure 1 is that ancestral SBW25 activates cellulose production in static broth culture and that polymer production allows cells to break through the meniscus and remarkably, grow transiently as micro-colonies on the surface. In the absence of cellulose production, cells are unable to penetrate the ALI and fail to reap the growth advantage that comes from a plentiful supply of oxygen (Fig. 2). Just how cellulose enables bacteria to break through the ALI is unclear. One possibility is that the polymer changes viscosity and this alone is sufficient to propel bacteria through the meniscus, another possibility is that the polymer alters surface charge and that altered electrostatic properties of the cells affects interactions with the surface (40).

Also unknown is the signal(s) that lead(s) to activation of cellulose production. What is clear is that known DGC-encoding pathways are necessary to transduce effects through to cellulose production. The fact that three pathways all contribute to differing extents points to complexity in the mapping between DGCs and the cellulosic target (41). It is tempting to suggest that the signal is oxygen, but this seems unlikely because it is incompatible with the previous finding that SBW25 and a cellulose defective mutant are equally fit in an oxygen-replete environment (24). Our suspicion is that the signal stems from some physical attribute of the ALI, possibly surface tension and Marangoni forces arising as a consequence of evaporation or production of surfactant – a subject that received momentary attention almost a century ago (42,43, 44).

The ecological significance of the behaviour is unknown. Assuming our observations are relevant to the natural environment and not just to laboratory culture, then one possibility is that cells use cellulose to colonise the ALI of water films on plant roots / leaves (the natural environment of SBW25 (45)) and use this environment to aid rapid and unimpeded dispersal. An additional benefit may then accrue on drying when the dispersed bacteria are bought back in contact with a solid substrate. Suggestive though that the growth extending up and out of the liquid surface may hint at a more complex and as yet unrecognised behaviour is the involvement of three DGC-encoding pathways. Why involve three pathways to regulate cellulose production when one would seem to suffice?

As colonisation of the surface begins to saturate, the heavier material on top becomes unstable and collapses in plumes typical of Rayleigh-Taylor instability. That such behaviour occurs is consistent with the thesis that cellulose is produced just at the meniscus and is not evenly distributed throughout the broth phase. Numerous consequences arise from the ensuing bioconvection, one of which is the rapid transport of water-soluble products. Our particular attention has been the fluorescent molecule pyoverdin, which by virtue of association with cells, is rapidly mixed from the point of production (the ALI) through the entire broth phase of the microcosm. Bioconvection additionally alters the chemical status of the environment, not only through mixing of extant products, but also through effects wrought by enhanced transport of oxygen.

Transport of pyoverdin has particular significance in light of a previous analysis of SBW25 populations propagated in static KB culture of an extended period (23, 46). Common mutant types that rose to prominence harboured mutations that abolished pyoverdin production. The evolutionary advantage of these mutants stemmed not from scavenging of pyoverdin (akin to “cheating”), but from avoidance of the cost of producing pyoverdin when it was not required (23). That pyoverdin is not required in the broth phase (because lack of oxygen means iron exists in the soluble ferrous state) is evident from the time-lapse movies (supplementary movie 7) where pyoverdin production is initiated exclusively at the ALI. However upon reaching the point of Rayleigh-Taylor instability, bioconvection due to cellulose rapidly transports pyoverdin into the broth phase where, in complex with iron, it serves to positively activate transcription of pyoverdin synthetic genes (47)- even though pyoverdin is not required by broth-colonising cells.

Imaging of cultures as reported here draws attention to the complexity and interdependence of biological, chemical and physical processes. A primary goal of the modelling exercise was to see just how far physical descriptions of measured phenomena such as plume velocity, bacterial density and pyoverdin concentration could account for observed dynamics. Similar approaches have been taken previously in analysis of microbial systems (48, 49). Specifically, our model shows how dynamical processes occurring in the liquid can be affected by biofilm formation at the ALI. It also reveals how proliferation of biomass affects the production and transport of pyoverdin. Additionally it accounts for physical transport of water-soluble products and the relative contributions of diffusion versus bioconvection to this process.

The model generates results consistent with cellulose production at the ALI being sufficient to generate Rayleigh-Taylor instability and initiate fluid movement. The specific mechanisms in the model involve the imbalance between the force of mass repartition in the fluid and the damping force of viscosity. The model also supports hypotheses concerning the critical role of cellulose in bioconvection: numerical resolution of the model showed plumes to have a velocity of ~1000μm·min^−1^ as observed in the experiment. Integrity of plumes – often tens of millimetres in length – is also explained by the model, and arises from the fact diffusion is a minor contributor to fluid dynamics relative to the effects of bioconvection. A further insight concerns ability of bioconvection to mix oxygen into the top few millimetres of the broth phase at a rate that is greater than its consumption. All these effects follow from the Rayleigh-Taylor instability wrought by the production of cellulose at the meniscus.

Together this study has shed new light on the role of cellulose — a widespread microbial product (50) — in colonisation of the ALI. Previous work has drawn attention to cellulose as an adhesive substance affecting the relationship between bacteria and solid surfaces (51, 52, 53,19), but these findings stem from study systems that do not provide opportunity for ALI colonisation and perhaps by design even select mutants that over-express cellulose and thus mislead as to ecological significance. This stated, cellulose may play different ecological roles in different organisms and under different conditions. Nonetheless, recognition that production of a polymer can modify an environment thus significantly changing the relationship between the organism and its environment – and the environment in a more general sense – has implications for understanding a range of environments and processes affected by ALI biofilms, such as those encountered in sewage treatment plants, marine and fresh water systems, and in terrestrial environments where transient films of moisture exist in soil pores and on plant surfaces. It also raises intriguing possibilities for future research on the importance of surface tension as a cue eliciting phenotypic responses in bacteria.

## MATERIALS AND METHODS

### Bacterial strain and growth conditions

The ancestral strain of *P. fluorescens* SBW25 was isolated from the leaf of a sugar beet plant at the University of Oxford farm (Wytham, Oxford, U.K.; (54)). The Δ*wssA-J* strain is deleted of the entire *wssA-J* operon (PFLU0300-PFLU0309) in the ancestral background and comes from (28). The Δ*wsp*, Δ*aws* and Δ *mws* were previously constructed by a two-step allelic exchange strategy (27).

Strains are cultured in King’s Medium B (KB) (55) at 28°C. KB contains (per litre) 20g bactoTM proteose peptone No. 3 (BD ref211693), 10 g glycerol, 1.5 g K_2_HPO_4_ and 1.5g MgSO_4_·7H_2_O.

To follow bacterial dynamics in experimental flasks bacteria were pre-cultured in KB overnight, centrifuged (6000rpm/3743rcf, 4min) and resuspended in fresh KB. The OD of suspended cultures was adjusted to an OD600nm of 0.8 and stored in 20μl aliquots containing 10μl of cultures of OD 0.8 and 10μl of 60v/v% autoclaved glycerol. The aliquots were conserved at −80°C.

To establish each experiment, a rectangular flask (Easy Flask 25cm^2^ Nunc) was filled with 20ml of KB medium. 20μl stock culture at −80°C was then thawed and inoculated in the KB at a final dilution of approx 10^4^cfu·ml^−1^. The flask was positioned in the setup shown in Figure 1 and incubated at 28°C in an IGS60 HERATHERM static incubator.

### Experimental setup to measure the dynamics of unshaken bacterial culture

The setup was designed and built to perform custom measurements and details are available from the authors upon request. The device comprises a laser-photodiode alignment to measure optical density in the flask and three cameras to observe the ALI as well as the biomass and the pyoverdin in the liquid phase.

To measure the optical density of the liquid phase a vertical profile was obtained by scanning with a laser-photodiode detector mounted on a lifter. A plastic piece that was produced by a 3D-printer on a stratasys fortus250 in ABS (yellow on Fig. 1) joined the laser-photodiode to a carriage that was free to slide on a vertical rail (ingus TS-01-15/TW-01-15) driven by a M10 cage bolt coupled to a M10 threaded rod. This ensured a precise vertical and horizontal positioning of the laser-photodiode alignment. The thread rode was smoothly rotated using a 7.2Vcc motor. A L293D power switch controlled by an Arduino board MEGA 2560 directed rotation of the motor. The thread rod rotation angle was measured with an optical encoder HEDS5500 500CPR. Ultimately, this allowed measurement of the vertical position of the laser beam with a resolution of ~3μm.

The photodiode was from Thorlabs (FDS1010), the laser a HLM1230 of wavelength 650 nm and power 5mW. To ensure that the laser did not harm bacteria the light was attenuated by a NE520B (Thorlabs) neutral density filter of OD=2. The optical density of the culture was evaluated by measuring the photo-current produced by the laser hitting the photodiode after it went through the flask. To ensure linearity between the intensity of light hitting the photodiode and its conversion in photo-current, the photodiode was polarised in inverse with 5V provided by a LM4040 electronic component. The photocurrent was estimated by measuring the voltage of a 437±5% kΩ resistor mounted in serial with the photodiode. The voltage was monitored by the Arduino MEGA 2560 board encoding a 0-5V analogic input on 10 bits giving a resolution of 5mV. After subsequent calibration, the signal acquired by the system allowed estimation of the bacterial concentration in the flask within a range of 10^7^ – 5 · 10^9^cell ·ml^−1^.

Synchronized with the laser-photodiode, were three cameras: a uEyeLE USB2.0 Camera, a 1/2” CMOS Monochrome Sensor, and a 1280×1024 Pixel equipped with a CMFA0420ND 4 mm ½ inch lens. The first camera (Fig. 1) records a bright field image of the vertical side view of the culture medium. The second camera is equipped with a band pass optical filter 470 ±10nm (Thorlabs FB470-10). It takes a side view of the culture medium. During its acquisition a 405nm laser (405MD-5-10-1235) illuminates the flask to excite pyoverdin fluorescence. The third camera takes a bright field image of the ALI with an angle of −45°. Acquisition of optical density data and photos were synchronized using a master script written in Python that recorded the data produced by the Arduino board and saved the photos taken by the cameras.

### Colonization of the air-liquid interface ALI (Fig. 4)

To observe the effect of Δwsp, Δ*aws* and Δ*mws* mutations on ALI colonization, we used 6-well plates Greiner bio-one 657160 filled with 8ml of KB. Each well was inoculated from glycerol stocks. The 6-well plates ere incubated at 28°C without shaking. Pictures were taken with a Nikon D7000 camera equipped with an AF-S DX NIKKOR 18-105mm f/3.5-5.6G ED VR objective.

### The advection-diffusion-convection model

The model uses six fields to describe the system: the vector field of the fluid vorticity (*ω*), the scalar field of the fluid stream function (Ψ) and the scalar fields of bacterial (*b*), oxygen (*o*), cellulose (*c*) and pyoverdin (*p*) concentrations. We also use a derived vector field that represents the velocity of the fluid (*u*). The model is valid for a three dimensional space but we estimate its validity in a two dimensional space in order to reduce the time of numerical computation. That is why we choose a fluid description in term of vorticity (*ω*) and the stream function (Ψ). This description gives two advantages for the numerical resolution of the model. First, the equation of the fluid incompressibility is solved by construction; second, the calculation of the vorticity vector can be reduced to the calculation of a simple scalar field (for more detail see (34)).

With the six fields given above come six partial differential equations that describe their dynamics.

The first equations related to the stream function. This scalar field is calculated by a Poisson equation:

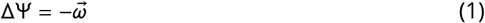

Where, Δ is the Laplace operator. The second equation deals with the vorticity reduced to a simple scalar field. Its dynamics can be derived from the Navier-Stokes (NS) equation:

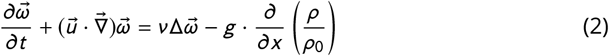

The left side of NS is the Lagrangian derivative of the vorticity. The right side contains damping of the vorticity by the viscosity *ν*, and a gravity term traduces the generation of vorticity due to the uneven spatial repartition of the mass density *ρ* relative to the density of the fluid medium *ρ*_0_. The operator 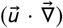 stem for the convective derivative. In the equation, the local mass densityρ takes into account the mass density of the liquid medium *ρ*_0_, bacteria *ρ_b_* and cellulose *ρ_c_*. Hence we consider the local mass density (*ρ*) as the sum of the mass contribution of the liquid medium, the bacteria and the cellulose. Explicitly the notation *ρ* stem for:

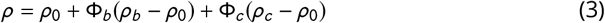

where Φ_*b*_ is the local volume fraction of the bacteria and Φ_*c*_ the local volume fraction of the cellulose.

To calculate the dynamics of the concentration of bacteria (*b*), cellulose (*c*) and pyoverdin (*p*), we write a diffusion-reaction-convection equation.

To write the third equation dealing with bacteria we make several assumptions:

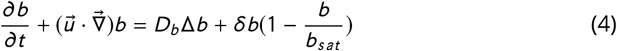

bacteria grow exponentially until they reach the saturation *b_sat_* measured experimentally (Fig. 2) and, bacteria consume oxygen that is dissolved into the liquid.

The left-hand side of the bacterial equation is the Lagrangian derivative applied to b. The right-hand side contains a diffusive term that takes into account the random motility of bacteria with a diffusion coefficient Db, and an exponential growth term with a rate *δ* that goes to zero when the concentration reach the *b_sat_* value.

The fourth equation describes the dynamics of the oxygen (*o*) field. Bacteria consume the oxygen at a rate γ.

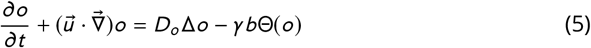

The coefficient of diffusion is Do. Oxygen consumption goes to zero when there is no more oxygen. This is ensure that multiplying the consumption term by the Heaviside function (Θ(*o*)) is 1 when o is above zero, but zero otherwise.

The fifth equation assumes that the cellulose is produced with an exponential rate (*α*) as long as the oxygen concentration is higher than *o*\* and the concentration of bacteria is higher than *b*\*. Additionally, cellulose production saturates when c tends to 1, its maximal value. The equation is:

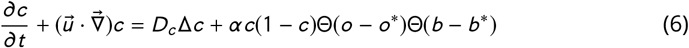

The sixth equation describes the dynamics of pyoverdin production. Provided that the local concentration of oxygen is sufficiently high bacteria produce pyoverdin according to the autocatalytic synthesis measured experimentally (Fig. 5D) with a rate *β*. The equation is:

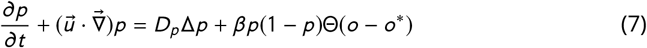

Here, pyoverdin production goes to zero when oxygen concentration is below *o*\* by multiplying the production term by a Heaviside function Θ(*o* – *o*\*).

Finally, to calculate the fluid velocity we use the derivative of the stream function where *u_x_* and *u_y_* stand for the horizontal and vertical components of the fluid velocity (*u*):

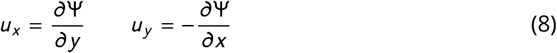

### Numerical Simulations

We used a finite difference method to solve the coupled reaction-diffusion-convection equations (56). The simulation was performed on a Linux system: Debian 4.9.51-1, gcc 6.3.0. The hardware CPU was an Intel(R)Core(TM) i7-7700K @ 4.2Ghz with 16GB RAM. The parameters used in the simulation displayed in Figure 7 are listed in Table 1.

**TABLE 1.**
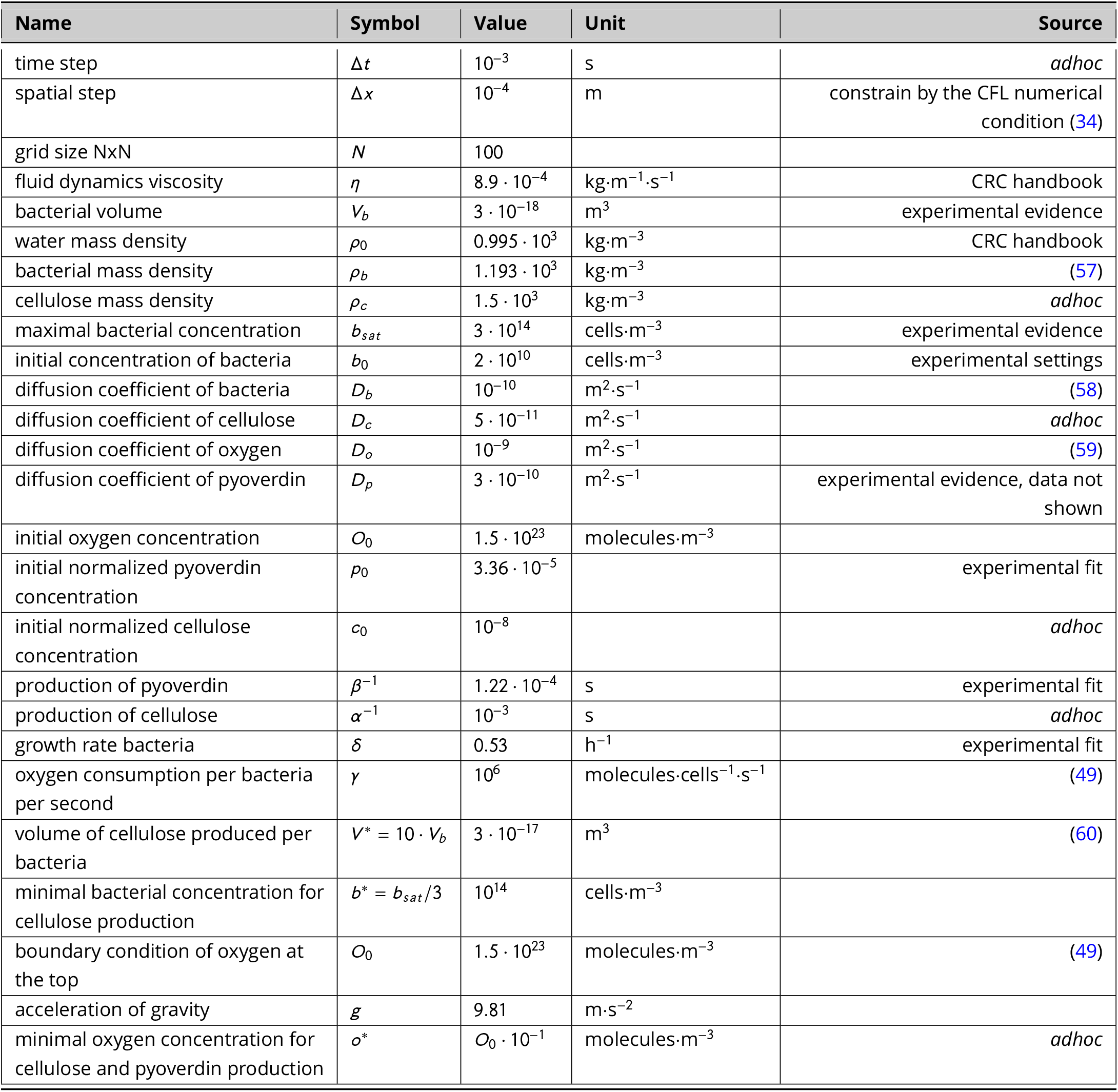
Parameters, values and sources of data used for the model.

## ACKNOWLEDGMENTS

We thank Nicolas Desprat, Clara Moreno Fenoll, Steven Quistad and Guilhem Doulcier for discussion and comment. MA was supported by HFSP grant RGP0010/2015.

## SUPPLEMENTAL MATERIAL

Supplementary movie file 1.

Name of file: supplementaryMovie1_ancestor_Surface.avi Growth of ancestral SBW25 imaged from camera 2.

Available at: https://figshare.com/projects/Causes_and_consequences_of_cellulose_production_by_Pseudomonas_fluorescens_SBW25_at_the_air-liquid_interface/59630

Supplementary movie file 2.

Name of file: supplementaryMovie2_Wss_Surface.avi Growth of SBW25 Δ*wssA-J* imaged from camera 2.

Available at: https://figshare.com/projects/Causes_and_consequences_of_cellulose_production_by_Pseudomonas_fluorescens_SBW25_at_the_air-liquid_interface/59630

Supplementary movie file 3.

Name of file: supplementaryMovie3_ancestor_Turbidity.avi Growth of ancestral SBW25 imaged from camera 1.

Available at: https://figshare.com/projects/Causes_and_consequences_of_cellulose_production_by_Pseudomonas_fluorescens_SBW25_at_the_air-liquid_interface/59630

Supplementary movie file 4.

Name of file: supplementaryMovie4_Wss_Turbidity.avi Growth of Δ*wssA-J* SBW25 imaged from camera 1.

Available at: https://figshare.com/projects/Causes_and_consequences_of_cellulose_production_by_Pseudomonas_fluorescens_SBW25_at_the_air-liquid_interface/59630

Supplementary movie file 5.

Name of file: supplementaryMovie5_ancestor_ALI.avi Growth of ancestral SBW25 imaged by SLR camera mounted directly above the well.

Available at: https://figshare.com/projects/Causes_and_consequences_of_cellulose_production_by_Pseudomonas_fluorescens_SBW25_at_the_air-liquid_interface/59630

Supplementary movie file 6.

Name of file: supplementaryMovie6_ancestor_Bioconvection.avi Replicate of ancestral SBW25 growth imaged by camera 1 with a high frequency frame rate.

Available at: https://figshare.com/projects/Causes_and_consequences_of_cellulose_production_by_Pseudomonas_fluorescens_SBW25_at_the_air-liquid_interface/59630

Supplementary movie file 7.

Name of file: supplementaryMovie7_ancestor_Pvd.avi Production of pyoverdin by the ancestral SBW25 imaged from camera 2.

Available at: https://figshare.com/projects/Causes_and_consequences_of_cellulose_production_by_Pseudomonas_fluorescens_SBW25_at_the_air-liquid_interface/59630

Supplementary movie file 8.

Name of file: supplementaryMovie8_pvd.avi Simulation of the pyoverdin dynamics.

Available at: https://figshare.com/projects/Causes_and_consequences_of_cellulose_production_by_Pseudomonas_fluorescens_SBW25_at_the_air-liquid_interface/59630

Supplementary movie file 9.

Name of file: supplementaryMovie9_velocity.avi Simulation of the velocity dynamics.

Available at: https://figshare.com/projects/Causes_and_consequences_of_cellulose_production_by_Pseudomonas_fluorescens_SBW25_at_the_air-liquid_interface/59630

Supplementary movie file 10.

Name of file: supplementaryMovie10_biomass.avi Simulation of the biomass dynamics.

https://figshare.com/projects/Causes_and_consequences_of_cellulose_production_by_Pseudomonas_fluorescens_SBW25_at_the_air-liquid_interface/59630

Supplementary movie file 11.

Name of file: supplementaryMovie11_O2.avi Simulation of the oxygen dynamics.

Available at: https://figshare.com/projects/Causes_and_consequences_of_cellulose_production_by_Pseudomonas_fluorescens_SBW25_at_the_air-liquid_interface/59630

Supplementary file 12.

Name of file: supplementaryFile12 Robustness of the simulations with respect to the parameters: *b*\*, *c*_0_, *γ*, *ρ_c_*, *V*\* and *o*\*.

Available at: https://figshare.com/projects/Causes_and_consequences_of_cellulose_production_by_Pseudomonas_fluorescens_SBW25_at_the_air-liquid_interface/59630

